# c-Abl Kinase Targets Tight Junction Protein ZO-2 in Regulation of Cell Migration and Morphology

**DOI:** 10.1101/2024.08.25.609487

**Authors:** Doo Eun Choi, Bomi Gweon, Jacob Notbohm, Xue-song Liu, Jeffrey J. Fredberg

## Abstract

c-Abl is a non-receptor tyrosine kinase involved in the regulation of cell migration and morphogenesis, but underlying mechanism remains unclear. Here, we report the identification of tight junction protein ZO-2 as a bona fide substrate of c-Abl. We show that c-Abl directly binds to and phosphorylates the C-terminus of ZO-2. In addition, c-Abl stimulates the activity of JAK1, which subsequently phosphorylates the N-terminus of ZO-2. Using the RNAi-mediated knockdown/rescue strategy, we demonstrate that c-Abl regulates cellular morphology and migration through targeting ZO-2 for phosphorylation. c-Abl activity is also associated with decreased traction forces exerted on the cell substrate and reduced monolayer tension, thus corroborating c-Abl kinase activity-mediated inhibition of cell migration. Collectively, our data uncover ZO-2 as a novel mediator for c-Abl-dependent regulation of cell migration.

## Introduction

Cell migration is a critical process in development, immune response, and maintenance of homeostasis. The Abelson tyrosine kinase (c-Abl) is a non-receptor tyrosine kinase involved in various cellular processes including cell migration, morphogenesis, proliferation, survival ^1^ and central nervous system development ^2^. Cytoplasmic c-Abl interacts with F-actin and G-actin to regulate cell migration and morphology changes through actin cytoskeletal reorganization ^3^. During the early stages of cell spreading, integrins stimulate c-Abl kinase activity ^4^, which subsequently interacts with focal adhesion proteins ^5,6^ and phosphorylate cytoskeletal regulatory proteins ^7–9^. The precise mechanisms by which c-Abl contributes to cell motility and morphogenesis, however, remains poorly understood.

In normal cells, the catalytic activity of c-Abl tyrosine kinases is tightly regulated as a critical component in various cellular responses ^10,11^. c-Abl can shuttle between the nucleus and the cytoplasm, facilitated by three nuclear localizing signals and a single nuclear exporting signal ^12,13^. c-Abl performs distinct functions dependent on its subcellular distribution: nuclear c-Abl is associated with G1 cell cycle arrest ^14^, DNA damage response ^15^, apoptosis ^16^, DNA repair and RNA polymerase II activation ^17^. In contrast, less is known about the cytoplasmic functions of c-Abl. Deregulated c-Abl kinase activity is implicated in abnormal motility associated with diverse pathological consequences including chronic myeloid leukemia (CML) ^18^, acute lymphocytic leukemia (ALL) ^19^ and neurodegenerative diseases such as Alzheimer’s ^20^ and Parkinson’s diseases ^21^. c-Abl knockout in mice results in embryonic or postnatal lethality, morphological abnormalities, T and B cell lymphopenia ^22,23^ and cardiac abnormalities ^24^.

Tight junction proteins ZO (Zonula Occludens, TJPZ)-1, ZO-2, and ZO-3 are phosphoproteins and belong to the MAGUK family. ZO proteins are multidomain proteins that recruit multiple and diverse molecules at the tight junction (TJ) region. Deregulation of ZO-2 function is associated with conditions such as family hypercholanemia ^25^, nonsyndromic progressive hearing loss ^26^ and various epithelial cancers including breast cancer ^27^, testicular carcinoma ^28^ and pancreatic duct adenocarcinomas ^29^. The loss or mutation of ZO proteins can result in the disruption of cell polarity ^30^ and tissue architecture ^31^.

In this study, we have identified ZO-2 as a novel substrate of c-Abl, which directly binds to and phosphorylates the ZO-2 protein. Multiple lines of evidence are provided to demonstrate that c-Abl-mediated ZO-2 phosphorylation plays a critical role in regulating cellular morphology and migration. We believe that these findings will enhance our understanding of the cytoplasmic functions of c-Abl and its key role in cell motility and morphogenesis, providing new insights into the molecular mechanisms underlying these processes.

## Results

### c-Abl Kinase Activity Induces Cell Morphology Changes

To examine the mechanism of c-Abl-mediated morphology and cytoskeleton change, we generated tetracycline-inducible c-Abl cell lines expressing either the kinase-active (KA) or kinase-inactive (KR) form of c-Abl. Western blot analysis confirmed that the expression of both c-Abl (KA) and c-Abl (KR) was markedly induced at 24 and 48 h after the addition of tetracycline (Tet) (Figure 1A). As expected, tyrosine kinase activity was detected only in c-Abl (KA) cells, as evidenced by the anti-Tyr immunoblot (Figure 1A) and was absent in c-Abl (KR) cells. Interestingly, there was a drastic change in cell morphology specifically associated with c-Abl kinase activity. When comparing c-Abl kinase-activated cells (KA + Tet) to the controls cells (KA) without Tet, a significant alteration of the cell monolayer was observed, resulting in island-like cells aggregates and a loss of typical single-layer epithelial cell morphology (Figure 1B). In contrast, c-Abl kinase-inactive cells (KR + Tet) showed no change in the morphology.

**Figure 1.**
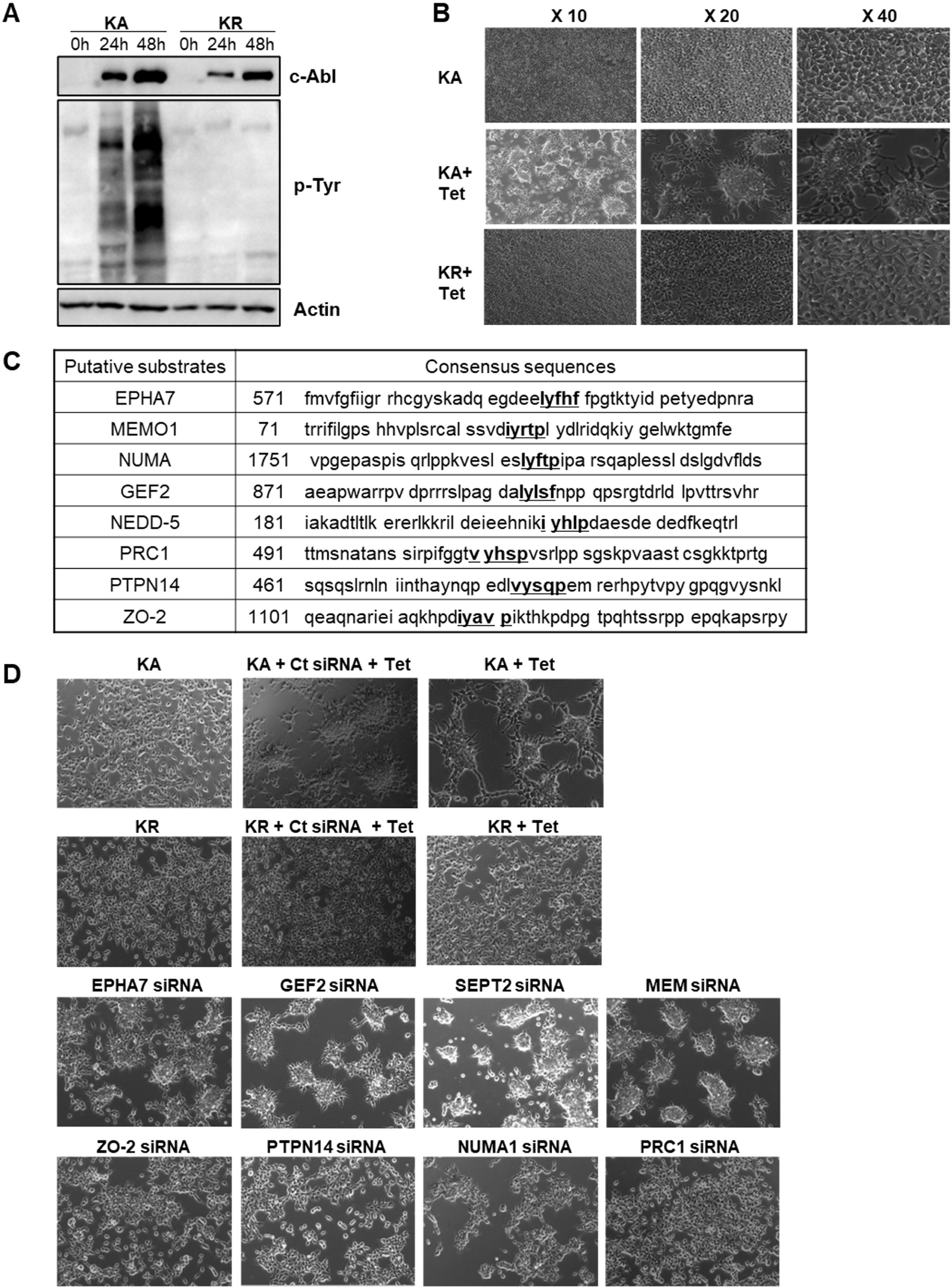
c-Abl induced-cell morphology change and potential substrates of c-Abl. (A) c-Abl-inducible cells, active c-Abl (KA) and inactive c-Abl (KR), were incubated for 24h and 48h after tetracycline treatment (1μg/ml). (B) c-Abl-inducible cell lines were induced using tetracycline to express KA and KR for 48 hours. (C) Eight putative candidates of c-Abl were selected by the consensus sequence of c-Abl. (D) Each of the 8 candidates was depleted by siRNA-mediated knockdown in KA cells with tetracycline. (Ct siRNA: control siRNA, KA: kinase active c-Abl, KR: kinase inactive c-Abl, + Tet: tetracycline was added).

Given that the observed morphological changes were dependent on c-Abl kinase activity, we hypothesized that c-Abl induces these changes by phosphorylating cellular proteins. We tested the hypothesis by identifying potential substrates of c-Abl with a focus on cytoskeletal proteins. In a proteomic analysis (Pull-down/MS), we detected tyrosine phosphorylation of 26 cytoskeletal proteins exclusively in cells expressing kinase-active (KA) form (Supplementary Table S1). From these, we selected eight candidates that contained the c-Abl phosphorylation consensus sequence (**I/V/L-Y-X-X-P/F)** ^32,33^ (Figure 1C).

We reasoned that the protein has to be expressed in order to mediate the effects of c-Abl. Using c-Abl activity-induced morphology change as functional readout, we assessed each of the eight candidates via siRNA-mediated knockdown. The result revealed that reduced expression of ZO-2, PTPN14 (Tyrosine-protein phosphatase non-receptor type 14), NUMA1 (Nuclear Mitotic Apparatus Protein 1), and PRC1 (Protein Regulator of Cytokinesis 1) significantly attenuated c-Abl activity-induced morphology changes (Figure 1D). In contrast, knockdown of EPHA7 (Ephrin Type-A Receptor 7), GEF2 (Rho/Rac Guanine Nucleotide Exchange Factor 2), SEPT2 (Septin-2) and MEMO1 (Mediator of Cell Motility 1) did not induce a similar attenuation and kept the aggregated morphology (Figure 1D). KR-expressing cells were included as controls to demonstrate that knockdown of each candidate gene alone did not have significant impact on cell morphology. To further confirm the involvement of ZO-2 in c-Abl-medicated cell morphology changes, binding experiments were performed.

### Tight junction protein ZO-2 is a novel substrate of c-Abl

To determine whether ZO-2 binds to c-Abl, we performed immunoprecipitation (IP). Lysates prepared from cells co-expressing Flag-ZO-2 and either c-Abl (KA) or c-Abl (KR) were subjected to anti-Flag IP. The result of immunoblot showed that ZO-2 binds to both c-Abl (KA) and c-Abl (KR), indicating that this binding is independent of kinase-activity. Consistent with the proteomic data, anti-p-Tyr immunoblot detected tyrosine phosphorylation of ZO-2 specifically in c-Abl (KA) expressing cells, but not in c-Abl (KR) expressing cells (Figure 2A).

**Figure 2.**
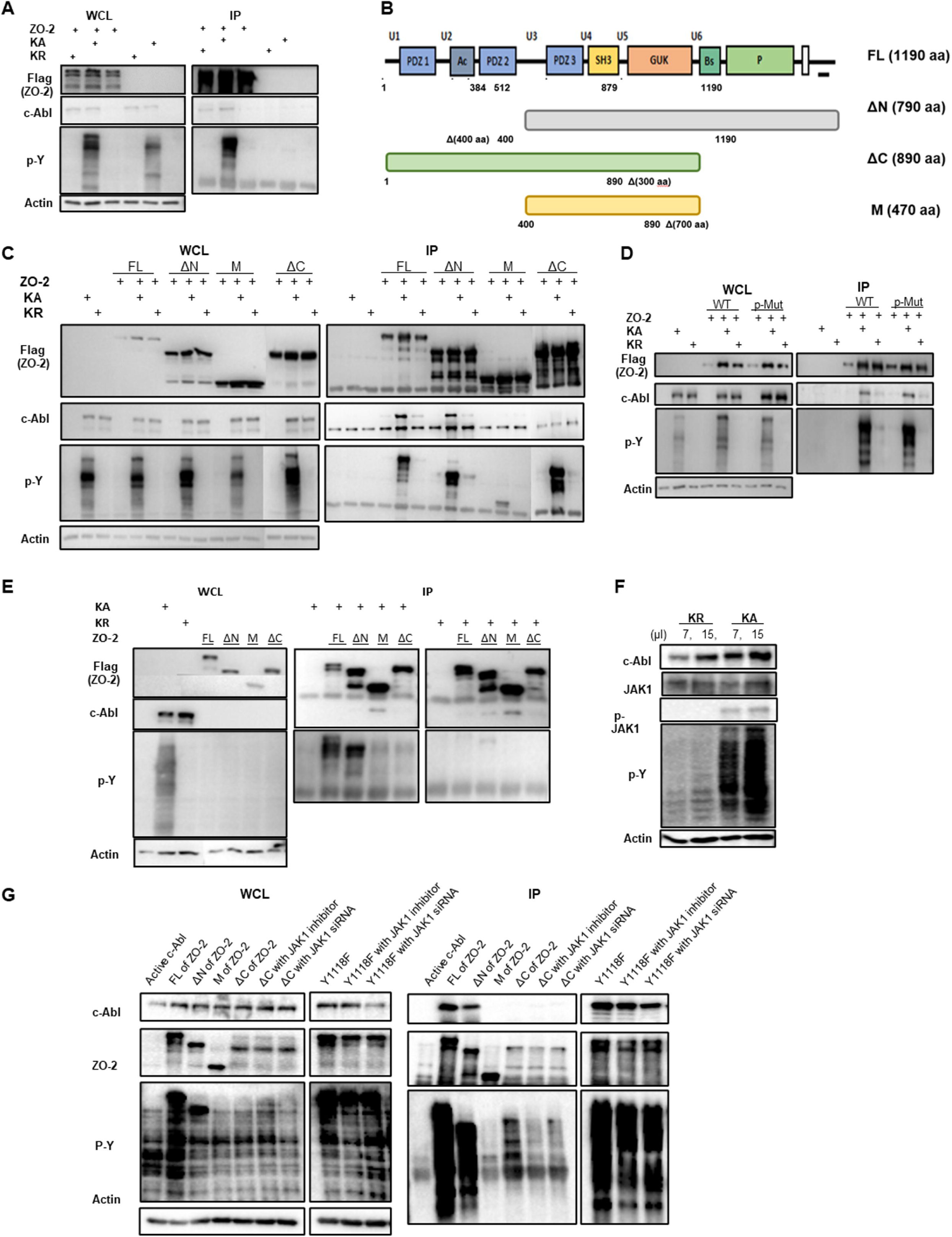
Tight junction protein ZO-2 as a new substrate of c-Abl. (A) Immunoprecipitation of ZO-2 and either KA (active) or KR (inactive) c-Abl. (B) Three truncated mutants. (FL: full length, ΔN: N-terminal deletion mutant, ΔC: C-terminal deletion mutant, M: middle part of ZO-2). (C) Either KA or KR was co-transfected with one of the four ZO-2 constructs (FL, ΔN, ΔC and M) into HEK 293 cells. Then, ZO-2 was pulled down with flag antibody (flag-ZO-2). (D) *In vivo* kinase assay with phospho-resistant mutant (Y1118F) and wild type of ZO-2. (E) *In vitro* kinase assay with FL and three deletion mutants of ZO-2. (F) Endogenous JAK1 and phosphorylated JAK1 were detected after induction of either KA or KR. (7 and 15 µ- amount of sample loaded.) (G) The tyrosine phosphorylation levels of ΔC mutant with JAK1 inhibitor and JAK1 siRNA.

To further characterize the interaction between c-Abl and ZO-2, we generated ZO-2 deletion mutants by dividing it into three fragments; N-terminal deleted (ΔN), only middle (M), and C-terminal deleted (ΔC) peptides (Figure 2B). IP-Western analysis revealed that c-Abl binds to both full length (FL) and ΔN fragments of ZO-2, but not to ΔC or the middle fragments (Figure 2C), indicating that c-Abl binds to the C-terminus of ZO-2.

Consistent with the location of c-Abl consensus phosphorylation site (Y1118) at the C-terminus, anti-p-Tyr detected strong phosphorylation of ΔN as well as FL ZO-2. Unexpectedly, ΔC-terminus, which had no detectable binding to c-Abl, was also found intensely phosphorylated by c-Abl (Figure 2C), suggesting the presence of intermediate tyrosine kinase mediating phosphorylation of ZO-2 N-terminus. In agreement with this possibility, the phospho-resistant (Y1118F) mutant of ZO-2 remained albeit slightly less phosphorylated (Figure 2D).

To ascertain the indirect phosphorylation of the ZO-2 N-terminus by c-Abl, we carried out an *in vitro* kinase assay. Purified full-length and deletion mutants of ZO-2 were incubated with either c-Abl (KA) or c-Abl (KR). Only the full-length and ΔN mutant of ZO-2, but not the ΔC mutant, were phosphorylated by c-Abl (KA), consistent with the notion that active c-Abl directly phosphorylates the C-terminal of ZO-2 and indirectly the N-terminus (Figure 2E).

Non-receptor tyrosine kinase JAK1 was reported to phosphorylate the N-terminus of ZO- 2 ^34^, and JAK1 is known to be activated by v-Abl ^35^, raising a possibility that JAK1 might be the intermediate kinases. To test this possibility, we first examined whether c-Abl could stimulate JAK1 activity using a specific phospho-JAK1 (Tyr1022/1023) antibody as a surrogate marker of JAK1 activation ^36^. Immunoblot analysis with this phospho-specific antibody detected JAK1 phosphorylation specifically in c-Abl (KA) but not c-Abl (KR) expressing cells, consistent with previous findings that c-Abl can activate JAK1 (Figure 2F) ^37^. We next investigated whether JAK1 is responsible for the phosphorylation of the N-terminus of ZO-2 using a combination of a JAK1 specific inhibitor and JAK1 siRNA. Phospho-tyrosine analysis indicated that inhibition of JAK1 was indeed associated with diminished phosphorylation of ZO-2 N-terminal peptides (Figure 2G). Collectively, these data support the conclusion that JAK1 acts as an intermediate kinase downstream of c-Abl to phosphorylate ZO-2 (Figure 2G).

### c-Abl-mediated ZO-2 phosphorylation plays significant role in cellular morphology

The results shown in Figure 2C, 2D, and 2E prompted us to identify additional phosphorylation sites on ZO-2 beyond Y1118. To this end, we performed phospho-proteomic analysis. Immuno- purified ZO-2 from either c-Abl (KA) or c-Abl (KR) expressing cells was subjected to MS/MS analysis of tyrosine phosphorylation. Indeed, apart from Y1118, we identified several additional sites of tyrosine phosphorylation in ZO-2 isolated from c-Abl (KA) but not c-Abl (KR) cells (Figure 3A).

**Figure 3.**
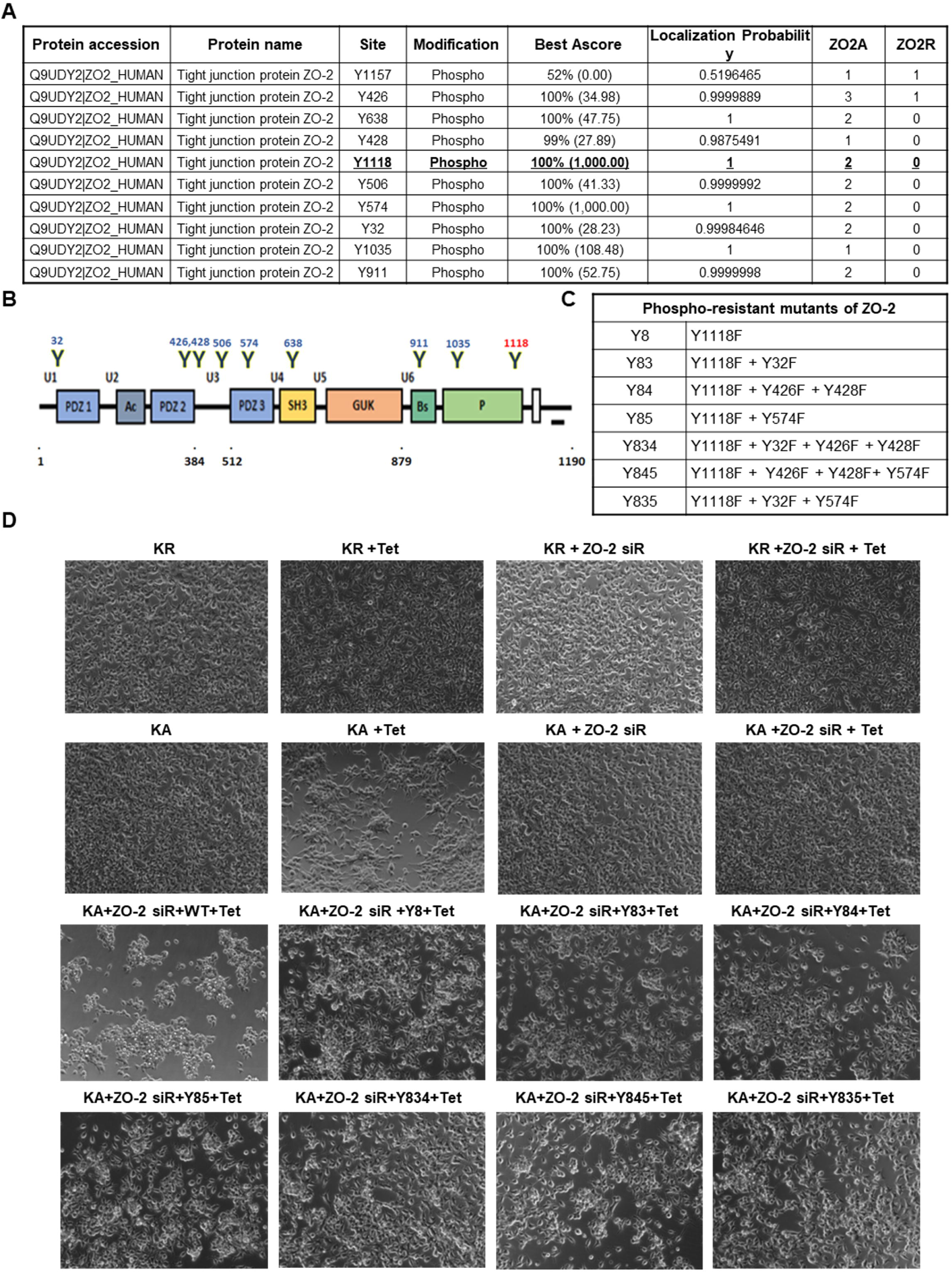
Cell morphology changes by c-Abl-mediated ZO-2 phosphorylation (A) Identification of tyrosine residues phosphorylated by c-Abl using Mass-Spectrometry analysis. (ZO2A: ZO-2 and KA were co-transfected into HEK 293 cells, ZO2R: ZO-2 and KR were co-transfected into HEK 293 cells). (B) Schematic of the ZO-2 structure and location of tyrosine phosphorylation sites. (C) List of the phospho-resistant mutants of ZO-2 and its abbreviations. (D) c-Abl-mediated cell morphology change was examined as a rescue experiment. As a control, either KA or KR cells was transfected with ZO-2 siRNA and treated with tetracycline. After siRNA transfection, either wild type (WT) or each of phospho-resistant mutants of ZO-2 was reintroduced into KA cells. Then, tetracycline (tet) was added to induce active c-Abl.

To determine the functional consequence of ZO-2 phosphorylation by c-Abl, we substituted each tyrosine (Y) residue with phenylalanine (F) individually or in combination (Figure 3B and 3C) and assessed each mutant for its ability to affect c-Abl activity-induced cell morphology changes. Specifically, we employed a depletion/rescue strategy to replace endogenous ZO-2 with each phospho-resistant mutant. To ensure the expression of exogenous ZO-2, we used ZO-2 siRNA targeting the 3’UTR of ZO-2 so that only the endogenous ZO-2 protein was knocked down. The results showed that re-expression of wild type ZO-2 in ZO-2- depleted cells almost completely restored c-Abl-induced morphology changes, indicating that exogenously expressed ZO-2 was able to functionally replace endogenous ZO-2 (Figure 3D). The Y1118F mutant of ZO-2 exhibited markedly reduced ability to rescue when compared with wild-type ZO-2, indicating the functional importance of Y1118 phosphorylation. Expression of other phospho-resistant mutants of ZO-2, with mutation sites in addition to Y1118F, in ZO-2-depleted cells resulted in even more compromised rescue, suggesting that these additional phosphorylation sites are also functionally important (Figure 3D).

### c-Abl-mediated ZO-2 phosphorylation plays a significant role in cell migration

To examine how c-Abl-induced ZO-2 phosphorylation affects cell migration, we employed a wound-healing assay that allows quantitative analysis. As expected, the control cells filled the wound or gap within 14 hours. In sharp contrast, the activation of c-Abl resulted in an almost complete halt of cell migration, leaving the gap largely unfilled (Figure 4A KA + Tet, and supplementary videos 1, 2, and 3). This effect appears to be kinase activity-dependent, as the expression of c-Abl (KR) under the same condition did not impair wound healing.

**Figure 4.**
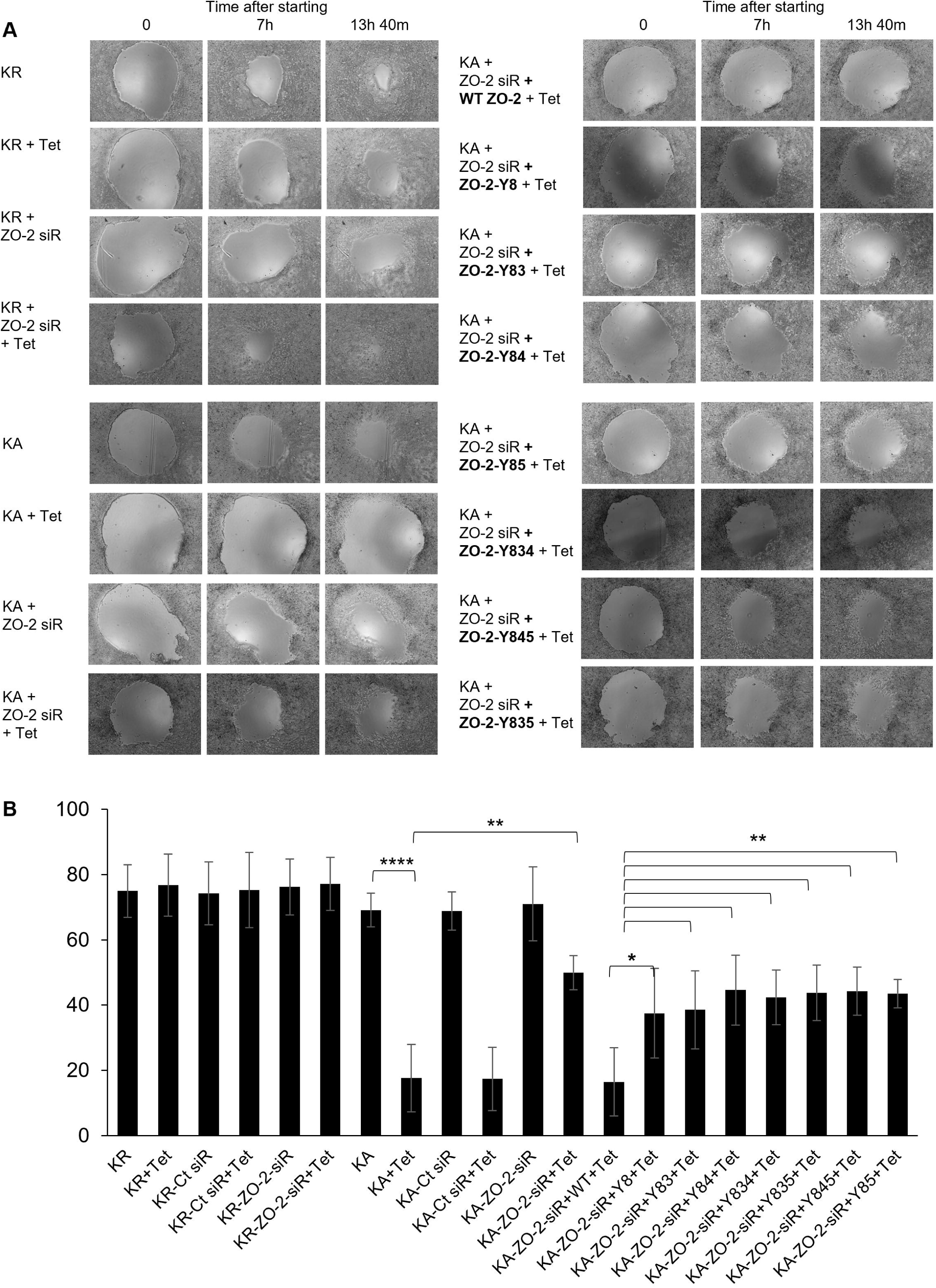
Inhibition of cell migration by c-Abl kinase activity and ZO-2 phosphorylation. (A) Wound healing assay. Tetracycline inducible KR cells and KA cells were used. Either wild type or each of phospho-resistant mutants was reintroduced into ZO-2 depleted KA cells and then c-Abl was induced using tetracycline. (Time-lapse imaging-10 min intervals, 0: starting point). (B) Quantification of the wound healing Assay. The ratio of the wound size (measured by area) before and after the wound closure. T-test was used for statistics.

We next investigated whether ZO-2 mediates c-Abl-induced inhibition of cell migration by employing a gene depletion/rescue strategy. Interestingly, c-Abl-induced inhibition of cell migration was almost completely lost upon siRNA-mediated depletion of ZO-2, as evidenced by the mostly filled gap (Figure 4A KA + Tet + ZO-2 siRNA, and supplementary videos 3, 4, and 5). Remarkably, the re-expression of wild-type ZO-2 in ZO-2-depleted cells fully restored the c-Abl-dependent inhibition of migration, demonstrating that ZO-2 is largely responsible for mediating this inhibitory effect on migration.

We then assessed the significance of ZO-2 phosphorylation in c-Abl-induced inhibition of cell migration by replacing wild-type ZO-2 (Figure 4A and Supplementary video 6) with various phospho-resistant mutants of ZO-2 (Figure 4A). The results confirmed the critical role of c-Abl-mediated ZO-2 phosphorylation in regulation of cell migration. Among the identified phosphorylation sites, Y1118 seemed to play a major role, while phosphorylation at additional sites was also contributed partially to the effect. This was demonstrated by comparing the ratio of the wound area at the start of the assay to the area of the same wound upon assay completion (Figure 4B).

### Tight junction protein ZO-2 plays an important role in c-Abl-mediated reduction of cell-ECM traction and cell-cell adhesion

As such, the wound-healing assay (Figure 4) revealed a significant role of c-Abl-mediated ZO-2 phosphorylation in cell migration. To understand how these changes in cellular morphology and motility following c-Abl activation relate to the adhesive functions of cells, we used EGTA (Ethylene glycol-bis (beta-aminoethyl)-N, N, N’, N’-t etraacetic acid) to challenge and disrupt cell-ECM and cell-cell adhesions. EGTA is well-known for inducing the loss of barrier function and increasing paracellular transport. Paracellular permeability across epithelial and endothelial cells is mainly regulated by TJs, which require optimal calcium concentration. Consequently, a decrease in extracellular calcium concentration leads to the disassembly of TJs ^38^. Therefore, EGTA, as a calcium chelator, is often used to assess TJ functions ^39,40^.

We employed 5 mM EGTA as the working concentration. As shown in Figure 5, eight conditions were examined: KR, KR + Tet, KR + ZO-2 siRNA, KR + ZO-2 siRNA + Tet, KA, KA + Tet, KA + ZO-2 siRNA, and KA + ZO-2 siRNA + Tet. Detachment of both cell-to-cell adhesion (indicated with red arrows) and cell-to-matrix adhesion (indicated with blue arrows) was observed in the time-lapse images. The results showed that dissociation in c-Abl kinase-inactive cells (KR, KR + Tet, KR + ZO-2 siRNA, and KR + ZO-2 siRNA + Tet) began approximately 120 seconds after the addition of EGTA. Interestingly, c-Abl activity markedly accelerated disassociation. Disruption of cell-cell adhesion in the c-Abl kinase-activated cells (KA + Tet) was observed at 65 seconds (Figure 5F). This early cell detachment in the c-Abl kinase-activated cells was attenuated by ZO-2 depletion (KA + Tet + ZO-2 siRNA) (Figure 5H), supporting our hypothesis that ZO-2 mediates the effect of c-Abl activation. In summary, these results indicate that c-Abl activation weakens both cell-cell and cell-matrix adhesion, and this functional effect is mediated by ZO-2.

**Figure 5.**
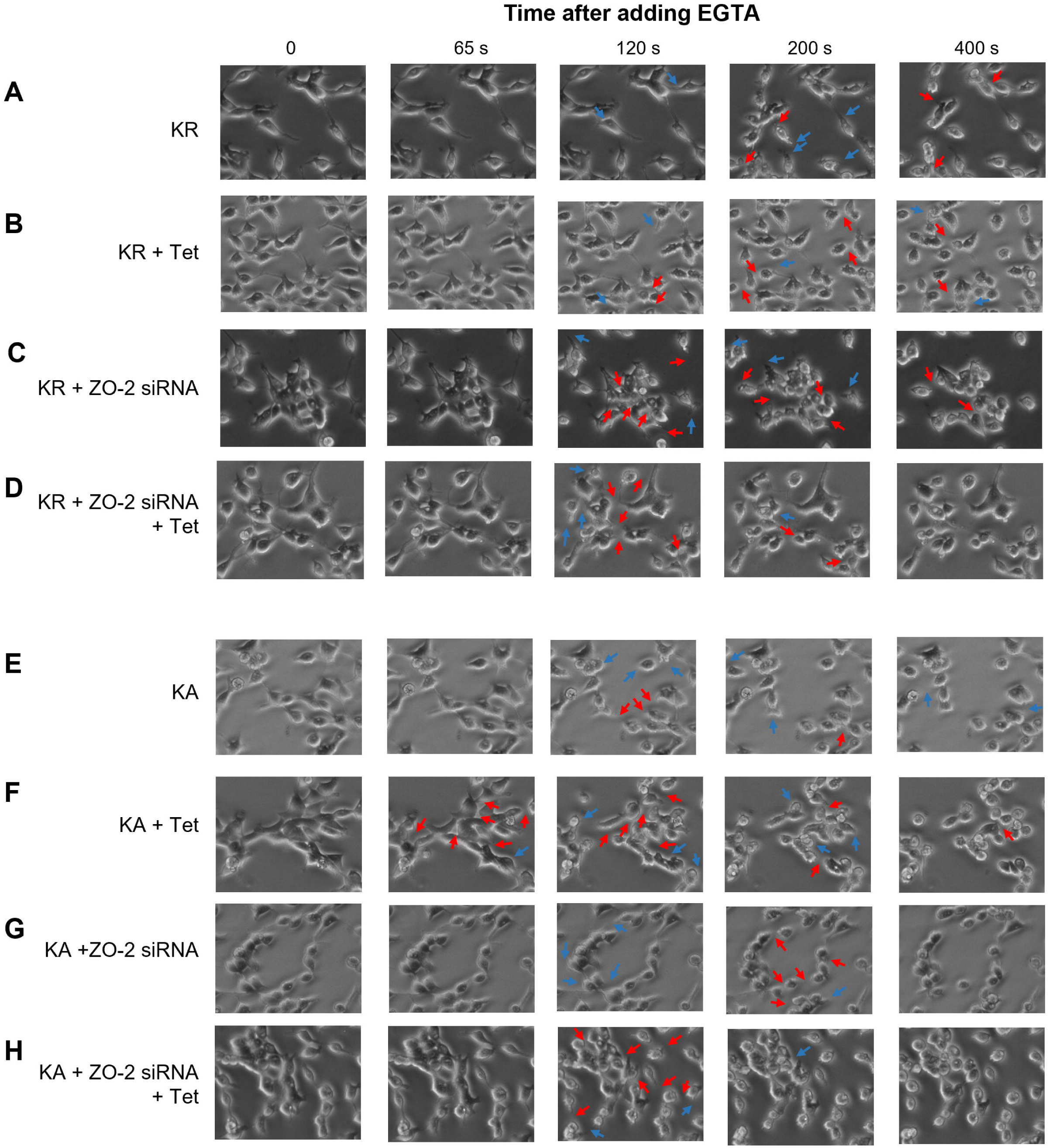
Early dissociation of cell interactions by c-Abl kinase activity and ZO-2. Early disruption of cell-cell and cell-substrate adhesion of KA + Tet cells (F) at 65 seconds. This phenotype was reversed by ZO-2 depletion (H). Disruption of cell-cell (red) and cell-matrix (blue) interactions is marked with arrows. Seconds indicates time after adding 5mM EGTA as a working concentration. Time-lapse images were captured every 2.5 seconds. (I) The schematic of c-Abl-induced cell behavior changes via ZO-2 tyrosine phosphorylation.

### Active c-Abl induces cell mechanical force changes across the cellular monolayer through ZO-2 tyrosine phosphorylation

Then, to precisely quantify the cell-to-matrix/substrate and cell-to-cell adhesion, we measured traction and tension within the cell monolayer. As depicted in Figure 6A, traction refers to the force per area exerted by cells onto the substrate. By employing traction force microscopy (TFM), the mechanical interactions between cells and the underlying matrix/substrate can be examined in a more direct and quantitative manner ^41–44^. We measured traction in cellular monolayers cultured under four selected conditions: KA, KA + Tet, KA + ZO-2 siRNA, and KA + ZO-2 siRNA + Tet. Representative traction maps are shown in Supplementary Figure S1. To compare traction quantitatively between c-Abl activated and non-activated cells, we plotted histograms showing the distribution of traction magnitude within the monolayer, calculated the median values from six to eight separate experiments, and then obtained the average of these median values (Figures 6B and 6C).

**Figure 6.**
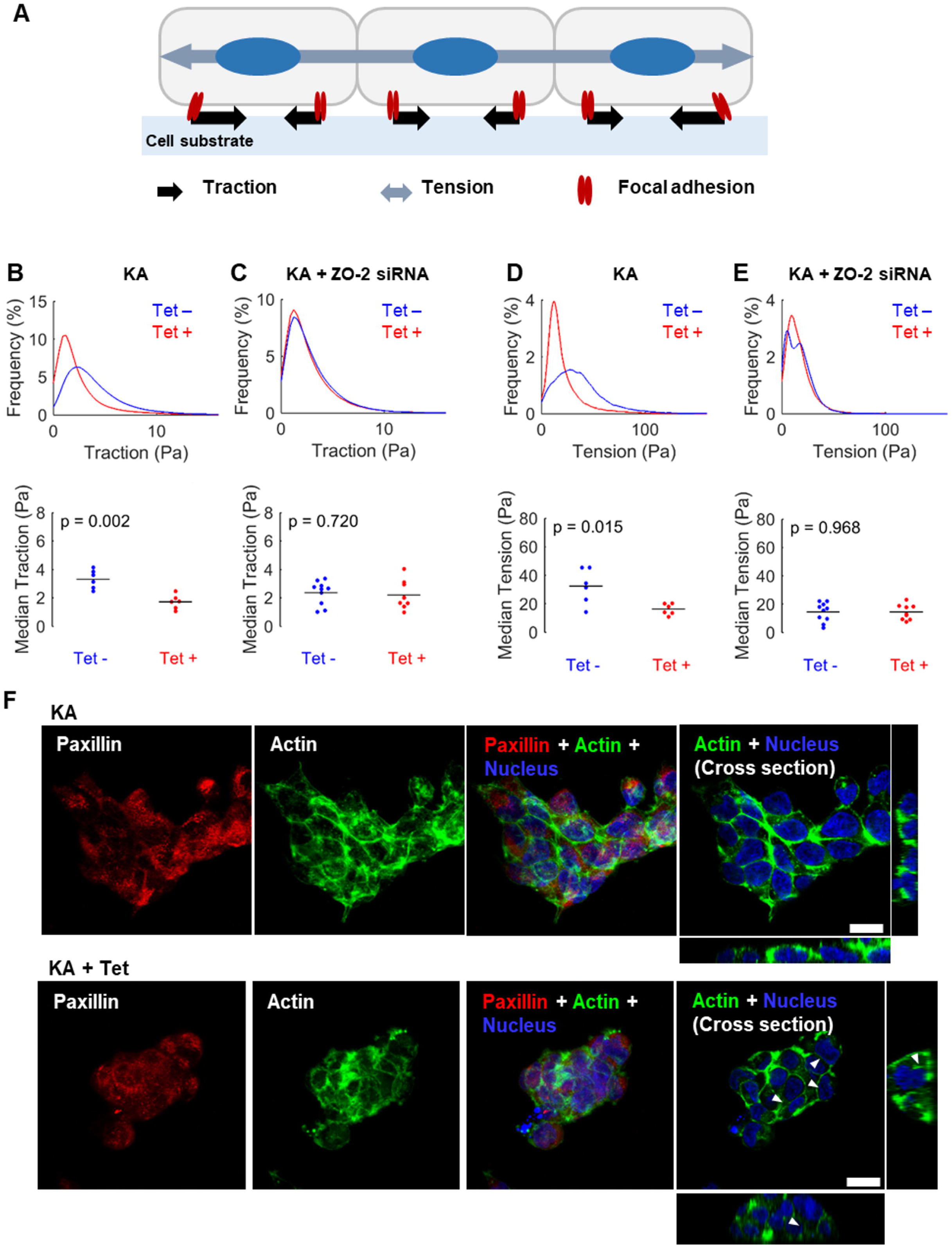
Reduced traction and tension by active c-Abl acting on ZO-2 and morphological changes within c-Abl induced cells. (A) Schematic representation of traction and tension within a group of cells. Black arrows indicate traction exerted on the substrate, and gray arrows represent tension across the cell monolayer. (B-E) Histograms show the relative frequency of cell-to-substrate traction (B, C) and cell-to-cell tension (D, E) in multiple different cell monolayers. The median traction and tension values for each monolayer are plotted as individual dots, with the mean value across experiments shown by a horizontal line (—) on each graph. (B, D): KA cells; (C, E): ZO-2-depleted KA cells; blue lines: without tetracycline (Tet -), and red lines: with tetracycline (Tet +). [n = 6 for KA cells and n = 8 for ZO-2-depleted KA cells, p-values from rank sum test] (F) Immunofluorescence image of KA cells with and without tetracycline. Paxillin (red), actin (green), and nucleus (blue) were stained. Scale bar = 100 μm.

As shown in Figure 6B, tractions exerted by c-Abl kinase-activated cells (KA + Tet) were significantly lower than those by KA cells without Tet, implying that active c-Abl perturbs and decreases the cell-to-substrate traction force. The force did not decrease by active c-Abl when ZO-2 was depleted by ZO-2 siRNA (Figure 6C), confirming that ZO-2 has a profound impact on c-Abl activity.

As a tight junction protein, ZO-2 is likely involved in the forces at the cell-cell junctions, specifically the tension across the cell monolayer that indirectly reflects monolayer integrity. As described in Figure 6A, tractions represent the forces transmitted to the substrate rather than those between neighboring cells. Therefore, to quantify intercellular forces within the monolayer, we employed monolayer stress microscopy (MSM) ^45,46^. MSM computes stresses (force per unit area) within the cell monolayer from the traction map by applying the principle of force equilibrium within the monolayer. From the data, we calculated the average principal stress within the cell monolayer, which we refer to as the tension. By assuming the cellular monolayer is a continuum, the computed tension reflects the overall trend of intercellular forces (Figure 6A).

Given that c-Abl kinase-activated cells (KA + Tet) led to reduced cell-cell adhesion under EGTA challenge conditions and decreased traction forces, it was anticipated that these cells would exhibit weaker monolayer tension compared to control cells (KA). Consistent with this expectation, our results revealed reduced tension within the c-Abl kinase-activated cellular monolayer (KA + Tet), measured at 16 Pa, compared to the control monolayer (KA), which was measured at 32 Pa (Figure 6D). In addition, the differences of tension mediated by active c-Abl were attenuated by ZO-2 depletion (Figure 6E).

Lastly, we investigated migratory machineries known to be responsible for contractile and adhesive forces in cells, such as actin and focal adhesion proteins. Paxillin, one of the major components of focal adhesions, did not show any significant changes in number or size due to c-Abl activity (Figure 6F). Actin stress fibers, however, became less prominent and thinner in c-Abl kinase-activated cells (KA + Tet) compared to control cells (KA) (Figure 6F). Since actin stress fibers are generally known to transmit contractile forces between cells and further between cells and substrate, these results align well with the reduced traction and tension observed.

## Discussion

c-Abl, a member of the non-receptor Src tyrosine kinase family, is known to contribute to signaling events that regulate diverse cellular functions. Depending on the localization of its substrates, c-Abl can influence both nuclear and cytoplasmic cellular processes. In this study, we focused on the cytoplasmic activity of c-Abl given that many cytoskeletal proteins were found to be tyrosine phosphorylated in response to c-Abl activation. Using cellular morphology as a functional readout to screen for c-Abl putative substrates, we identified ZO-2 as the protein chiefly responsible for c-Abl kinase-dependent regulation of cell morphology and migration (Figure 3 and 4).

Binding studies demonstrated a direct association of c-Abl to the C-terminus of ZO-2 in a kinase-independent manner. This binding allows c-Abl targeting ZO-2 for phosphorylation (Figure 2A and C). Specifically, c-Abl phosphorylates Y1118, a residue within the consensus sequence at ZO-2 C-terminus (Figure 2E). Apart from this direct phosphorylation site, c-Abl also induces phosphorylation at additional sites indirectly through other tyrosine kinases, including JAK1, as revealed by siRNA-mediated knockdown and the use of a specific inhibitor (Figure 2G). In agreement with the presence of additional phosphorylation sites, phospho-proteomic analysis (Pull-down/MS) uncovered eight new phosphorylation sites in the ZO-2 protein (Figure 3A). Collectively, these results indicate that c-Abl activity induces ZO-2 phosphorylation at multiple sites via a combination of direct and indirect mode of action.

A critical role of ZO-2 in mediating c-Abl activity-induced effects on cell morphology was investigated by depletion/rescue experiments. We first established the essential role of ZO-2 by showing that knockdown of ZO-2 abrogated, whereas the re-expression of ZO-2 restored c-Abl-dependent regulation of cell morphology and migration (Figure 3D and 4). In line with this kinase activity-dependent regulation, we showed that substituting wild-type ZO-2 with phospho-resistant mutants resulted in compromised rescue. Quantitative analysis revealed a major role of phosphorylation at Y1118, as well as other sites albeit to a lesser extent, in c-Abl activity-dependent regulation of cell migration.

The wound-healing assay also provided convincing evidence indicating that c-Abl kinase activity mediates the inhibition of cell migration. In line with our results, Kain and Klemke ^47^ have shown that c-Abl can reduce the rate of cell migration. Specifically, embryonic fibroblasts derived from *abl ^-/-^ arg ^-/-^* mice fill in a wounded area more rapidly than normal counterparts, whereas reintroduction of wild-type c-Abl into *abl ^-/-^ arg ^-/-^*cells with inhibits cell migration by preventing of Crk-p130 ^CAS^ (CAS) complex formation. Additionally, cell movement is enhanced by treatment with the Abl inhibitor STI571 and inhibited by overexpression of active c-Abl, reinforcing the notion that c-Abl plays a negative role in cell migration ^48^.

Further supporting this conclusion is the observation that c-Abl activity was associated with reduced traction and tension within the cell monolayer (Figure 6B to 6E). Cells generate contractile force provided by myosin-based motors, and this cellular force is transmitted to the matrix/substrate underneath them and cells next to them via the cytoskeleton and focal adhesions. The less distinct actin stress fibers in c-Abl kinase-activated cells (KA + Tet) (Figure 6F) are consistent with these findings, as these actin stress fibers are crucial for the transmission of contractile forces. Previous reports have shown that a certain level of contractile force is necessary for cells to gain thrust and migrate forward ^49, 50^. Therefore, the overall changes in cellular structure, traction, and tension observed in c-Abl kinase-activated cells (KA + Tet), as summarized in Figure 7A, likely contribute significantly to the reduced motility induced by c-Abl activity.

**Figure 7.**
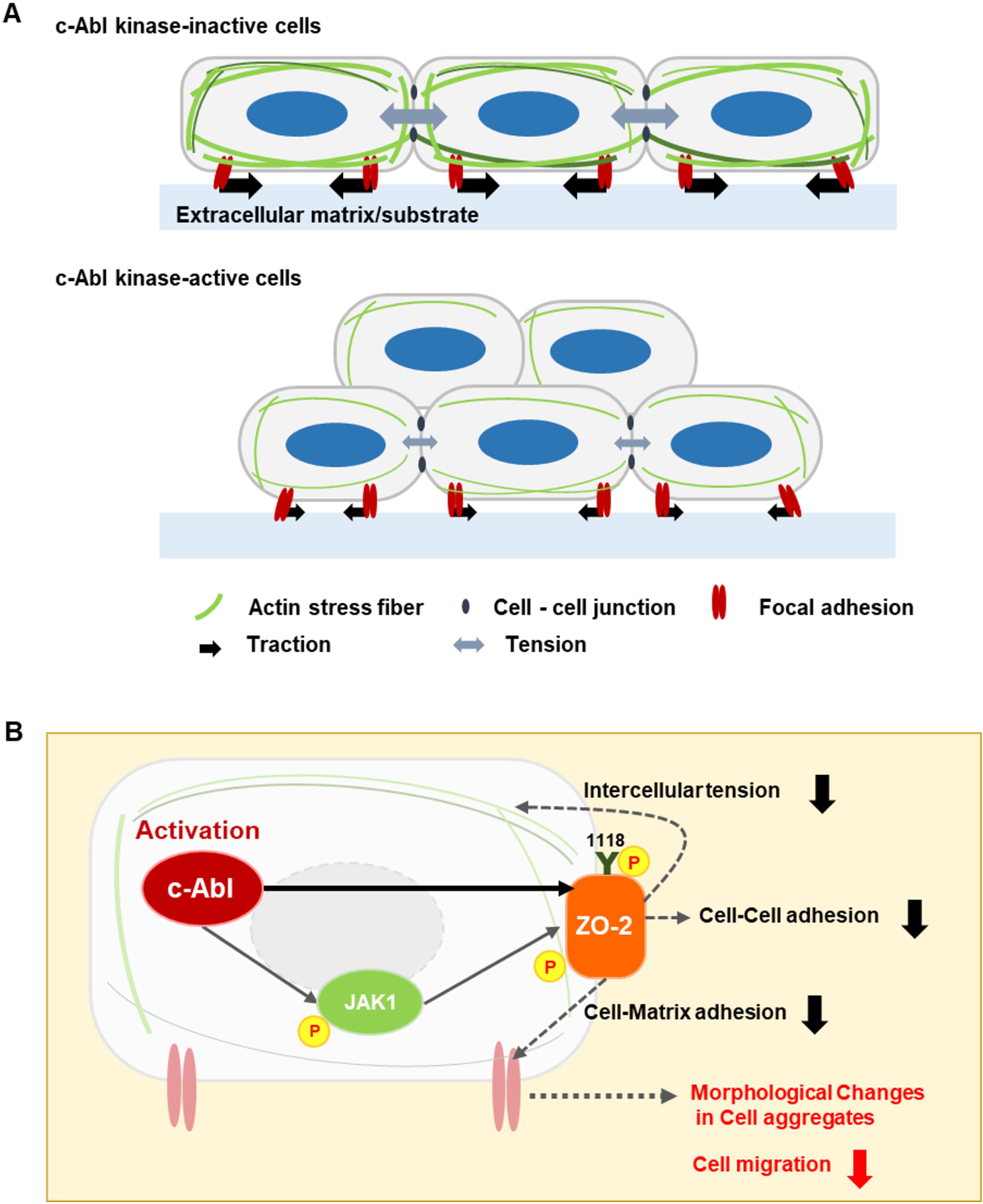
Schematics diagram of the changes within the cells by active c-Abl. (A) Illustrates the morphological and architectural changes in cells due to active c-Abl, and (B) depicts the signaling pathway activated by c-Abl. The activation of c-Abl leads to changes in intercellular tension, cell-cell adhesion, cell-matrix adhesion, and overall cell morphology.

ZO-2 has been implicated as an adaptor between actin and E-cadherin, contributing to the regulation of proteins important for cellular polarity ^51^. Hernandez et al ^52^, reported that ZO-2 silencing in epithelial cells disrupts the gate and fence functions of TJs, and alters the monolayer architecture ^53^. However, the impact of ZO-2 phosphorylation on cellular structural features and subsequent cellular functions has not been fully understood. Our study demonstrated that c-Abl-mediated ZO-2 phosphorylation contributes to c-Abl-induced alterations in cell morphology and migration, uncovering a novel mechanism of post-translational modification in functional regulation of ZO-2. Saito et al. have shown that ZO-2 can also be phosphorylated by the tyrosine kinase c-Src and interacts with its negative regulator, C-terminal Src kinase (Csk) ^54^. Additionally, tyrosine phosphorylation of ZO-1 and ZO-2 by v-Src has been shown to weaken junctional sealing ^55^. These findings align well with our results, demonstrating that ZO-2 phosphorylation through c-Abl activation (KA + Tet) induces a weakening of cell-cell adhesion (Figure 5) and decrease in monolayer tension (Figure 6D and 6E). Based on these results, we can also assume that phosphorylation of ZO-2 likely influences its interaction with other proteins, impacting its adaptor function and thus its role in regulating cellular polarity. Further studies will be necessary to test this possibility.

Given the aggregated morphological features and restricted migration in c-Abl activated cells (KA + Tet), one might anticipate stronger cell-cell adhesions. However, as shown above, our results clearly demonstrate that c-Abl activation and subsequent ZO-2 phosphorylation weaken, rather than strengthen, cell-cell adhesions. Therefore, the pronounced morphological changes leading to cellular aggregates in c-Abl activated cells appear to be induced through other mechanisms. Numerous prior studies have demonstrated that cellular aggregates can form under conditions of low cell-substrate adhesion. For instance, when seeded on a non-adhesive PolyHEMA, SCp2 (mouse mammary epithelial cells) and SCg6 (myoepithelial-like mouse-derived mammary cells) were shown to lose their attachment to the ECM/substrate and typical morphology, forming various patterns of aggregation ^56^. The presence of similar cellular clumps in c-Abl activated cells (Figure 1B) suggests that cell-substrate adhesion is dramatically decreased, indicating that c-Abl may play a key role in weakening cell adhesion to the substrate. In this context, we found that activation of c-Abl weaken cellular traction (mechanical interaction between the cell and the matrix/substrate).

This effect of c-Abl is mediated by ZO-2, revealing a novel function of ZO-2 in the regulation of cell adhesion and migration. ZO-2 has been reported to directly interact with junction adhesion molecule-A (JAM-A or JAM-1), which regulates cell adhesion and migration through dimerization by indirectly controlling β1 integrin level ^57^. We speculate that c-Abl-dependent phosphorylation of ZO-2 would affect its interaction with JAM-A and consequent cell adhesion and migration. Further studies are warranted to characterize the functional interaction between ZO-2 and JAM in the context of cell migration.

In conclusion, our data demonstrate that ZO-2 is a pivotal mediator of c-Abl-induced alterations in cell migration, adhesion, and morphology. The direct and indirect phosphorylation of ZO-2 by c-Abl leads to a reduction in both cell-cell and cell-matrix adhesion, which subsequently causes significant morphological alterations in cell aggregates, ultimately contributing to impaired cell migration (Figure 7B). The phosphorylation-mediated mechanism in functional regulation of ZO-2 may have important implications in various physiological and pathological contexts involving cell-cell interactions, cell attachments to the matrix, altered cell migration, and cellular morphological changes.

## Materials and Methods

### 1. Cell Culture, Transfection and c-Abl Induction

Human 293 cells were cultured in Dulbecco’s modified Eagle’s medium (DMEM) supplemented with 10% heat-inactivated fetal bovine serum (FBS), 100 μg/ml streptomycin and 100 units/ml penicillin and incubated at 37℃ in 5% CO2. Tet system approved FBS (Clontech) was used for 293FT-TREX inducible cell lines (active c-Abl KA and inactive c-Abl KR). For transfection assays, cells were seeded at least 12 hours before transfection. Cells were transfected with Lipofectamine 2000 reagent (Life Technologies). Lipofectamine 2000 reagent and plasmids were diluted in Opti-MEM medium (Sigma). For tetracycline induction, 293FT-TREX KA and KR inducible cell lines were treated with 1μg/ml of tetracycline for indicated time points to induce c-Abl expression.

### 2. Immunoprecipitation and Western Blot Assay

Harvested Cells were rinsed with PBS and lysed in lysis buffer (50 mM Tris pH 8.0, 150 mM NaCl, 5 mM EDTA, 0.5% NP-40, 2 mM PMSF, 20 mg/ml aprotinin, 25 mM NaF and 0.2 mM sodium orthovanadate). After centrifugation, supernatant was incubated with Anti-flag M2 magnetic beads (Sigma) for 4 hours at 4 °C. The samples were washed three times, and the supernatant were removed. 2 x SDS loading buffer was added and boiled. Prepared samples were loaded on a 12% SDS-PAGE gel. Gels were transferred to nitrocellulose membrane and incubated with blocking buffer, primary antibody, and secondary antibody (washed with TBS-T in between each incubation time). The blots were visualized by ECL. Antibodies used in Western blot analysis were c-Abl monoclonal antibody (K2, Santa Cruz), flag monoclonal antibody (Flag M2, Sigma), ZO-2 polyclonal antibody (Santa Cruz), phospho-tyrosine monoclonal antibody (4G10, Upstate Biotechnology), actin monoclonal antibody (Sigma), Jak1 polyclonal antibody (Cell signaling), and phospho-Jak1 (Tyr1022/1023) polyclonal antibody (Cell signaling).

### 3. Binding Assay and *In Vivo* Kinase Assay

Either c-Abl KA or KR were co-transfected with the ZO-2 into 293 cells. ZO-2 was isolated by immunoprecipitation (IP) using Anti-flag M2 magnetic beads (Sigma). Western blot analysis was performed with anti-c-Abl (K2, Santa Cruz), anti-flag (Flag M2, Sigma), anti-phosphotyrosine (4G10, Upstate Biotechnology) and anti-actin (Sigma) antibodies.

### 4. *In Vitro* Kinase Assay

ZO-2 was enriched with IP and incubated with either active c-Abl KA or inactive c-Abl KR in kinase buffer (25mM Tris pH 7.4, 10mM MgCl2, 1mM MnCl2, 0.5mM DDT and 10μM ATP) at 30°C for 15 min. The reaction was stopped by boiling in SDS sample buffer, and separated by SDS-PAGE. Western blot assay was used with anti-phosphotyrosine (4G10, Upstate Biotechnology), anti-c-Abl (K2, Santa Cruz), anti-flag (Flag M2, Sigma), and anti-actin (Sigma) antibodies.

### 5. Site-specific mutation

To generate phospho-resistant mutant plasmids of ZO-2 by substitution, PCR was performed with synthetic oligonucleotide primers using the QuikChange site-directed mutagenesis kit (Agilent, Stratagene). To digest the parental DNA template and to select for mutation-containing synthesized DNA, the mutant plasmids were treated with Dpn I endonuclease (target sequence: 5’-Gm6ATC-3’) which is specific for methylated and hemimethylated DNA. The nicked plasmids incorporating the desired mutations were then transformed into E. coli. The mutated plasmids were then be purified.

### 6. Immunofluorescence

Cells were fixed in 3.7% formaldehyde solution in PBS, permeabilized with TX-100 (0.01%), and incubated with blocking buffer (1% bovine serum albumin) and washed with PBS in between each incubation time. The samples were incubated with the indicated primary antibodies overnight at 4 °C and then, with all secondary antibodies included DAPI stain to visualize nuclei, mouse Alexa fluor 488 and rabbit Alexa fluor 647 conjugated antibodies. Cells were mounted and visualized using a fluorescence microscopy.

### 7. siRNA Transfection and Quantitative-real-time-PCR (q-RT-PCR) Analysis

Cells were transfected with siRNA using RNAiMAX reagent (Invitrogen) and medium was changed after 24 hours. Total RNA was isolated using Trizol/chloroform extraction as directed by manufacturer’s instructions (Invitrogen) and cDNA was generated. Quantitative PCR amplifications were performed using the QuantiTec SYBR Green PCR kit (Qiagen) and the Monitor Thermo cycler.

### 8. Mass-Spectrometry Analysis

Flag-ZO-2 was co-transfected with either c-Abl KA or KR. Cells were lysed in lysis buffer (50 mM Tris pH 8.0, 150 mM NaCl, 5 mM EDTA, 0.5% NP-40, 2 mM PMSF, 20 mg/ml aprotinin, 25 mM NaF and 0.2 mM sodium orthovanadate). ZO-2 was pulled downed with flag tag-conjugated beads and washed three times.

### 9. Optical Microscopy

Time lapse images of the cells were captured every 5 minutes using phase contrast on a DMI6000B microscope stand with a 5x NA 0.12 objective and a DFC345FX CCD camera (Leica). The imaging environment was maintained at 37°C/ 5% CO2 in a heated enclosure. For the EGTA assay, images were captured every 2.5 seconds.

### 10. Wound Healing Assay

For more accurate quantification of the wound closure rate circular wounds were introduced in 293FT KA/KR cell monolayers grown in a 96 well plate. To make a circular wound, we placed a polydimethylsiloxane (PDMS) pillar (diameter=0.5 mm) in each well before seeding the cells. The PDMS pillars were made by mixing the silicon elastomer and the cross-linker (Sylgard 184, Dow Corning) at 10:1 ratio and cured overnight on a hot plate at 80°C. Two c-Abl inducible lines (KA and KR) were plated and transfected with either control siRNA or ZO-2 siRNA. The cells were counted and plated in the prepared 96-well plates with each well having a PDMS pillar (Figure 3). The next day, either wild type or each plasmid of ZO-2 phospho-resistant mutants was transfected into ZO-2-depleted KA cells. Cells were incubated for 24∼30 hours. For active c-Abl induction, 1μg/ml of tetracycline was treated to the samples. 6 hours later, the pillar was removed leaving a cell-free area. Immediately after removing the PDMS pillar, phase contrast images of every wound in each well of the 96 well plate were captured every 10 minutes for 14 hours. We obtained images for various cells in parallel using the microscope’s motorized stage.

### 11. Traction Force Microscopy and Monolayer Stress Microscopy

Polyacrylamide gels were prepared as a substrate for computing cell tractions. Gels with Young’s modulus of 1.2 kPa were made by preparing a solution of 3% acrylamide (Bio-Rad), 0.11% bisacrylamide (Bio-Rad), 2 mg/mL acrylic acid N-hydroxysuccinimide ester (NHS; Sigma), 0.014% 0.5 µm fluorescent particles (Life Technologies) and 0.033% ammonium persulfate (Bio-Rad). 0.05% N,N,N’,N’-tetramethylethylenediamine (TEMED; Bio-Rad) was used to catalyze the reaction. 24 µL of the polyacrylamide solution was placed onto a glass-bottomed dish and covered with an 18 mm diameter coverslip. During polymerization, the gels were centrifuged upside down at 115 g for 8 min to move the fluorescent particles to the top surface of each gel. After centrifuging, the gels swelled in water overnight at 4°C. To grow cells on the gel, the gel surface was functionalized by adding 200 μl of 2 mM sulfosuccinimidyl 6 (4’- azido-2’-nitrophenyl-amino) hexanoate (sulfo-SANPAH; ProteoChem) to the surface of the gels and photoactivating with a UV lamp for 10 min. The gels were then rinsed and coated with 250 μl of 100 μg/ml collagen type l (Advanced BioMatrix) for about 8 hours at 4°C. After the surface treatment, gels were rinsed 3 times with phosphate buffered saline and warmed up to room temperature. Then, cells were trypsinized from the culture flask and seeded on the gel. A small amount (4 μl) of dense cell suspension (8x10^6^ cells/ml) was plated on each gel covered with pre-warmed culture medium, and the gels kept in the incubator for about 48 hours to allow cells to form a monolayer. To induce c-Abl, half of the monolayers were incubated with tetracycline for 12 hours (KA + Tet) and the other half was kept in tetracycline free medium (KA).

Time lapse images of cells (phase contrast) and particles beneath the polyacrylamide gel surface (fluorescence) were captured every 5 minutes for 6 hours using the Leica microscope described above. After the time lapse, the cell medium was removed, the cells were rinsed with phosphate buffered saline, and Trypsin was added. After waiting 10 minutes, the cells were gently aspirated to remove them from the polyacrylamide substrate, and a final image of the fluorescent particles was captured in the stress-free state. Polyacrylamide substrate displacements were computed using image correlation. Tractions applied by the cells to the substrate were computed from the displacements with Fourier transform traction microscopy ^58^, using an implementation for finite substrate thickness ^59,60^. Stresses between the cells were computed by applying a force balance to the traction data using monolayer stress microscopy ^61,62^. The in-plane stress tensor computed with monolayer stress microscopy has three unique components, two representing normal stresses and one representing shearing stresses. Since the shearing stresses are typically ≥ 4 times smaller than normal stresses, we report the average of the two normal stress components, which is a measure of the tension between neighboring cells.

### 12. EGTA Assay

5mM EGTA (Ethylene glycol-bis (beta-aminoethyl)-N, N, N’, N’-t etraacetic acid) was used as a working concentration. EGTA was added 30 seconds later after running of time lapse imaging. Time lapse images of cells (phase contrast) were captured every 2.5 seconds using the Leica microscope described above.

### 13. Statistical Analysis

Data are generally presented as mean values with standard deviations. Differences between two groups were assessed using the Student’s t-test. For the traction and tension analysis, data are presented as median values, and differences between groups were evaluated using the Rank-Sum test.

## Acknowledgement

The authors would like to thank Professor Zhi-Min Yuan for supervising, constructive discussion, and supporting this research.

## Author Contributions

Doo Eun Choi, Conceptualization, Methodology, Validation, Formal analysis, Investigation, Data curation, Writing – original draft, Writing – review and editing, Project administration; Bomi Gweon, Conceptualization, Methodology, Investigation, Writing – original draft, Writing – review and editing, Funding acquisition; Jacob Notbohm, Software, Validation, Data curation, Writing – review and editing; Xue-song Liu, Investigation, Methodology; Jeffrey J. Fredberg, Resources, Writing – review and editing

## Data Availability

The original data in this article will be shared upon reasonable request to the corresponding authors.

## Abbreviations

c-Abl, Abelson tyrosine kinase Tet, Tetracycline TJ, Tight Junction ZO, Zonula Occludens

