## Supplementary Table S1; Supplementary Figure S1 for "c-Abl Kinase Targets Tight Junction Protein ZO-2 in Regulation of Cell Migration and Morphology"

| Protein | Sequence source | Sequence number | presence of consensus sequence |
|---|---|---|---|
| Actin | NCBI | NP_001092.1 | PARTIAL |
| Activity-regulated cytoskeleton-associated protein (Arg3.1) | NCBI | NP_056008.1 | PARTIAL |
| Alpha-actinin 1 | GenBank | AAH15766.1 | PARTIAL |
| Catenin alpha-1 (CAP102) | GenBank | AAH00385.1 | PARTIAL |
| Dynamin binding protein | NCBI | NP_056036.1 | PARTIAL |
| Ephrin type A recepter 7 precursor (EPHA7) | Swiss-Prot | Q15375.3 | FULL |
| Four and a half LIM domains protein (FHL-1) | UniProtKB/Swiss-Prot | Q13642.4 | PARTIAL |
| Protein MEMO 1 (Mediator of ErbB2-driven cell motility) | NCBI | NP_057039.1 | FULL |
| Myosin-10 (heavy chain, nonmuscle) | NCBI | NP_005955.1 | PARTIAL |
| Nuclear mitotic apparatus protein 1 (NUMA1) | NCBI | NP_006176.2 | FULL |
| Partitioning-defective 3 homolog (PARD-3) (ASIP) ephrin-interacting protein (PHIP) | Swiss-Prot | Q8TEW0.2 | PARTIAL |
| Plastin-2 (LCP-1) | NCBI | NP_002289.2 | PARTIAL |
| PDZ and LIM domain protein 1 (Elfin) | NCBI | NP_066272.1 | PARTIAL |
| PDZ and LIM domain protein 5 (Enigma homolog) | NCBI | NP_006448.3 | PARTIAL |
| Profilin-1 | NCBI | NP_005013.1 | PARTIAL |
| Protein phosphatase 1 regulatory subunit 12A (myosin phosphatase target subunit 1) | GenBank | BAA22378.**1** | PARTIAL |
| Protein regulator of Cytokinesis 1 (PRC1) | Swiss-Prot | O43663.2 | FULL |
| Rho/Rac guanine nucleotide exchange factor 2 (GEF2) | NCBI | NP_001155855.1 | FULL |
| Septin-2 (protein NEDD5) | NCBI | NP_006146.1 | FULL |
| Src substrate cortactin (Amplaxin) | NCBI | NP_005222.2 | PARTIAL |
| Tyrosine-protein phosphatase non-receptor type 14 (Pez) (PTPN14) | NCBI | NP_005392.2 | FULL |
| Tight junction protein ZO-2 (Zonula Occludens 2 protein) | NCBI | NP_004808.**2** | FULL |
| Tubulin alpha-1A chain | UniProtKB/Swiss-Prot | Q71U36.1 | FULL |
| Tubulin beta-2 chain | UniProtKB/Swiss-Prot | P68371.1 | PARTIAL |
| Tubulin alpha-2 | GenBank | ABD72607.1 | FULL |
| Vimentin | NCBI | NP_003371.2 | PARTIAL |

**Table S1** A total of 26 cytoskeletal proteins were identified as tyrosine-phosphorylated exclusively in cells expressing the kinase-active (KA) form, as determined by pull-down/MS proteomic analysis. Among them, 10 contain the c-Abl phosphorylation consensus sequence.

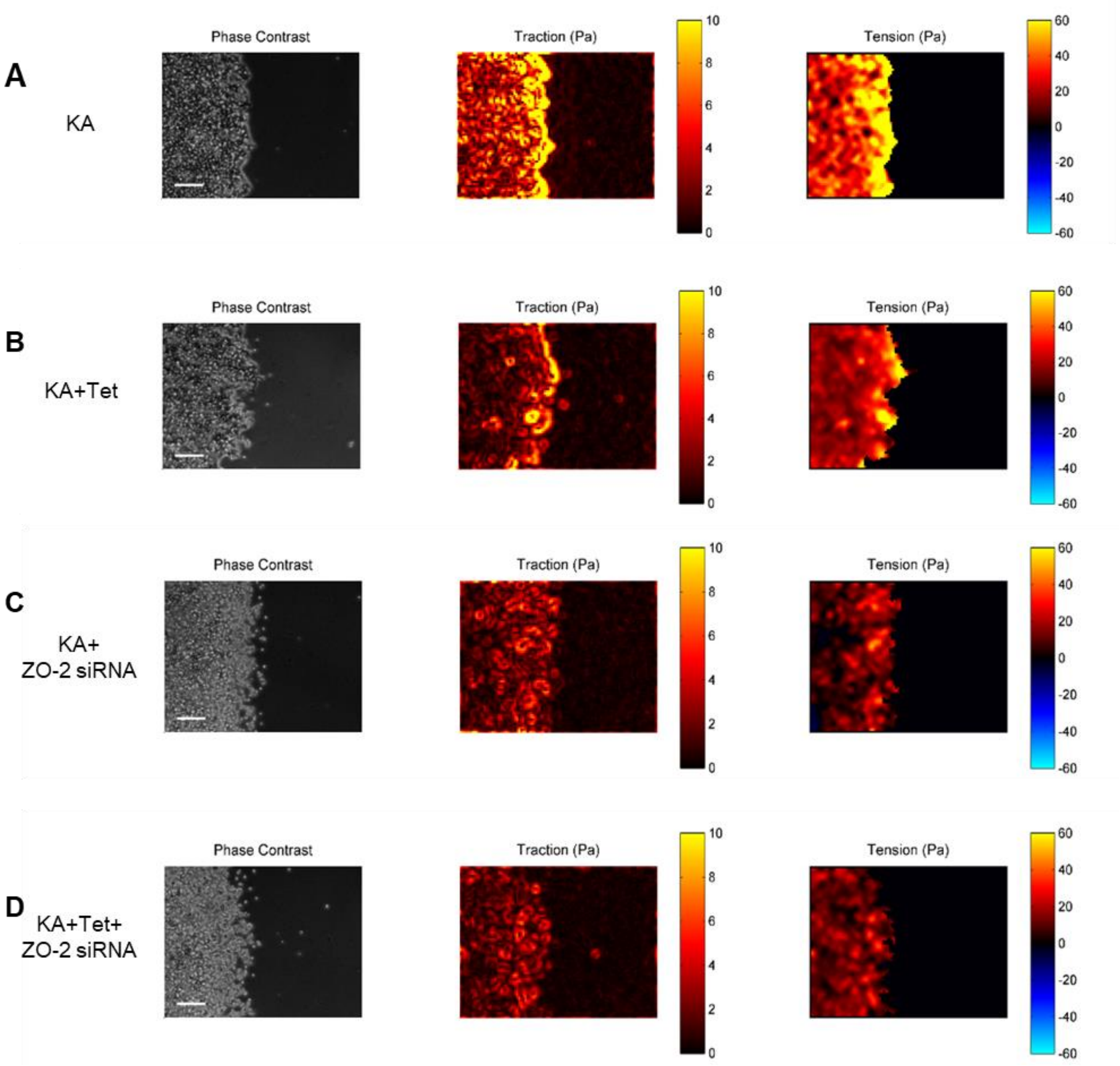


**Figure S1** Representative phase contrast images along with corresponding traction (Pa) and intercellular tension (Pa) maps. The images depict four different conditions: (A) KA, (B) KA + Tet, (C) KA + ZO-2 siRNA, and (D) KA + Tet + ZO-2 siRNA.

**Video S1** Wound healing process in cells expressing the kinase-inactive (KR) form of c-Abl.

**Video S2** Wound healing process in cells expressing the kinase-inactive (KR) form of c-Abl with Tet treatment.

**Video S3** Wound healing process in cells expressing the kinase-active (KA) form of c-Abl.

**Video S4** Wound healing process in cells expressing the kinase-active (KA) form of c-Abl with Tet treatment.

**Video S5** Wound healing process of cells expressing kinase-active (KA) form of c-Abl, treated with ZO-2 siRNA and Tet

**Video S6** Wound healing process of cells expressing kinase-active (KA) form of c-Abl, treated with ZO-2 siRNA, followed by transfection with WT ZO-2 and Tet treatment.
